# A hitchhiker’s guide to Europe: mapping human-mediated dispersal of the invasive Japanese beetle

**DOI:** 10.1101/2023.10.06.561184

**Authors:** Leyli Borner, Davide Martinetti, Sylvain Poggi

## Abstract

Early detection of hitchhiking pests requires the identification of strategic entry points via transport. We propose a framework for achieving this in Europe using the Japanese beetle (*Popillia japonica*) as a case study. Human-mediated dispersal has been responsible for its introduction into several continents over the last century, including Europe where it is listed as a priority pest. Furthermore, interceptions far from the infested area confirm the risk of unintentional transport within continental Europe. Here, we analyse how three modes of transport - air, rail and road - connect the infested area to the rest of Europe. We ranked all European regions from most to least reachable from the infested area. We identified border regions and distant major cities that are readily reachable and observed differences between modes. We propose a composite reachability index combining the three modes, which provides a valuable tool for designing a continental surveillance strategy and prioritising highly reachable regions, as demonstrated by recent interceptions.

**Significance statement:** Species can be moved long distances by unintentional human transport. Such events can lead to their introduction into non-native areas where they can cause biological invasions. Using the case study of the Japanese beetle, a hitchhiker invasive insect, we propose a framework for identifying entry points for human-transported pests within Europe. We consider how the European infested area is connected to the rest of the continent by three modes of transport: air, rail and road. We propose a methodology that considers the three modes to identify potential entry points. This framework could assist authorities in designing surveillance strategies to achieve early detection of pests in Europe.

## 1. Introduction

The increasing global movement of goods and people provides countless opportunities for pests to move around the world (1). Insects can hitchhike on a variety of modes of transport, including planes, ships, trains and trucks, facilitating their entry into new regions and, if conditions are suitable, subsequent establishment. Understanding the role of transport in the spread of insect pests is therefore crucial to developing effective surveillance strategies (2).

The Japanese beetle (*Popillia japonica*) is a prime example of a hitchhiker pest. Native to Japan, it was accidentally introduced into the United States of America at the beginning of the last century, causing a major invasion that still persists today. From there, it was introduced to the Azores archipelago in the 1970s, and more recently to continental Europe. After its first detection in Italy in 2014, the beetle has spread to an area of more than 16,500 km^2^ covering parts of northern Italy and southern Switzerland (3). Due to its potential impact on the environment, food safety and economic balances, it has been listed as a priority quarantine pest by EU authorities (4). Furthermore, several interceptions of the beetle far from the infested area have raised concerns about possible introductions to other parts of Europe.

In this paper, we present a novel approach to map the potential human-mediated dispersal of the beetle from the infested area to the rest of Europe. We considered three transport networks - air, rail and road - that are relevant to the beetle’s pathways of entry from the infested area. We looked at how reachability, i.e. the likelihood of entry from the infested area, varied between transport modes. Finally, we used a composite reachability index to identify the most likely points of entry and used interceptions sites to test our reachability maps.

## Materials & Methods

We first assessed the extent of the infested area, taking into account both the municipalities where the presence of the beetle was confirmed and the neighbouring municipalities included in the buffer zone, according to 2022 official reports (5). As there are no seaport in this area, the three major modes of transport that can facilitate the spread of the beetle were considered: air, rail and road. We used available data on movements of commercial flights, commercial trains, and freight trucks to represent the three modes of transport. For air and rail transports, we considered the number of passengers (resp. trains) departing from airports (resp. stations) in the infested area and arriving at airports (resp. stations) outside that area via a direct flight (resp. train), during the adult emergence period (May to August). For road freight, we used the estimated number of trucks departing from infested NUTS3 regions (EU Nomenclature of territorial units for statistics) and arriving in uninfested NUTS3 regions (6). Finally, we combined the information provided by the three transport modes using a Pareto front ranking method. This method classifies NUTS3 regions into ordered groups of decreasing composite reachability. Further details on data and methods can be found in the SI-extended methods. Data processing and analyses were performed using R version 4.2.1 (7).

## Results

The 2022 infested area covers approximately 16,500 km^2^, spanning over five regions and more than 1900 municipalities in Italy and Switzerland. Within this area, there are six airports and 540 train stations. Twenty NUTS3 regions are considered infested because they contain at least one infested municipality.

Reachability from the infested area vary between modes of transport (Figure 1A-C). Our analysis shows that of all NUTS3 regions directly reachable by at least one transport mode from the infested area, 7% and 10% are reachable by rail and air respectively. Meanwhile, almost all of these regions (99.8%) are reachable by road freight. From the infested area, 422 train stations in five countries, 160 airports in 30 countries and 1446 NUTS3 regions in 33 countries can be reached by commercial trains, planes, and freight trucks, respectively. With the exception of a few distant major EU cities, rail transport network mainly connects areas adjacent to the infested area and most of Italy and Switzerland. The road freight network is both local, with Switzerland, northern and central Italy highly reachable from the infested area; and international, with many major EU cities also highly reachable. Finally, the air transport network from the infested area is almost exclusively international, with direct access to all major cities in the EU.

**Figure 1:**
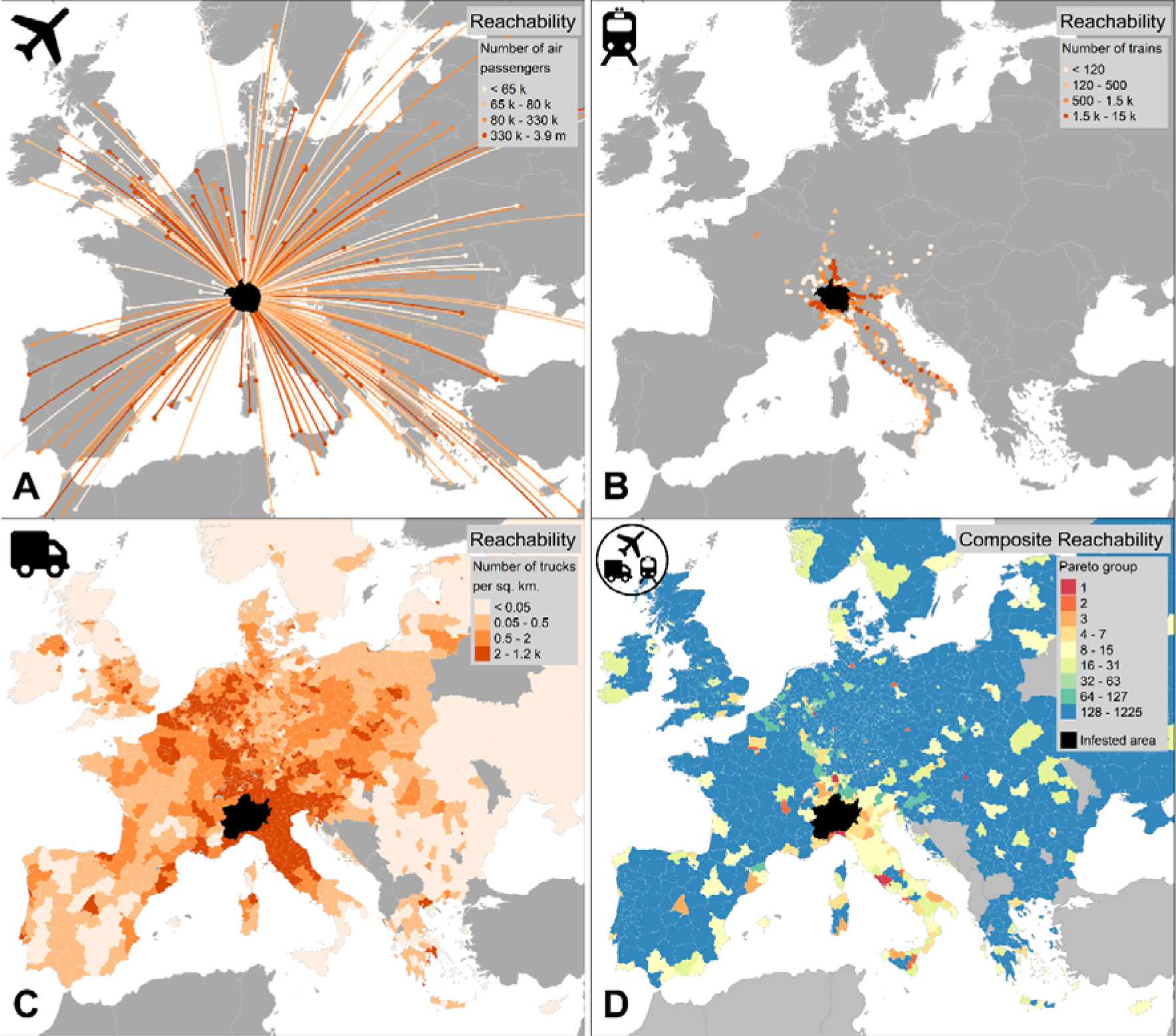
Reachability of Europe for *Popillia japonica* from the infested area (in black), by air, rail, road transport, and combined modes (composite index). Quantile-classified reachability maps showing, **A** the number of passengers arriving at airports, **B** the number of trains arriving at stations, and **C** the number of trucks per square kilometre reaching NUTS3 regions, departing from the infested area. Darker colours correspond to higher reachability. **D** Composite reachability map, i.e. likelihood of entry for NUTS3 regions grouped by Pareto fronts from most to least reachable. Warmer colours correspond to higher reachability.

Our composite reachability index, which combines air, rail and road transport, classifies NUTS3 regions into ordered groups, from most to least reachable (Figure 1D). Interestingly, 13 NUTS3 regions are reachable by all transport modes and belong to the five most reachable groups. However, not all 62 NUTS3 regions in these five groups are reachable by all modes. The distribution of highly reachable regions is scattered and anisotropic. Most of these regions are located far from the infested area, showing that reachability is not correlated with distance. Furthermore, most of the regions with medium and high reachability are located either in Italy and Switzerland (due to the predominant local and national nature of rail and road freight transport) or in major European cities (especially along an N-S longitudinal transect crossing France, Germany and Benelux).

## Discussion

We mapped the distribution of reachability of Europe by air, rail and road transport from the current Japanese beetle infested area. We found that reachability varies by mode, and detected topological features of transport networks, ranging from predominantly local and national (rail and road transport) to almost exclusively international (air). Our proposed composite reachability index highlighted a few scattered highly-reachable major cities across Europe, as well as a cluster of high reachability in Italy and Switzerland, which could be potential entry points.

Interestingly, all Japanese beetle interceptions that have occurred in Europe over the past 5 years have been reported in regions that our analysis identified as highly reachable (Figure 2), providing preliminary evidence that our approach is robust. Furthermore, an outbreak was detected in July 2023 in one of the nine regions we have identified as the most reachable from the infested area (Zurich region, Switzerland).

**Figure 2:**
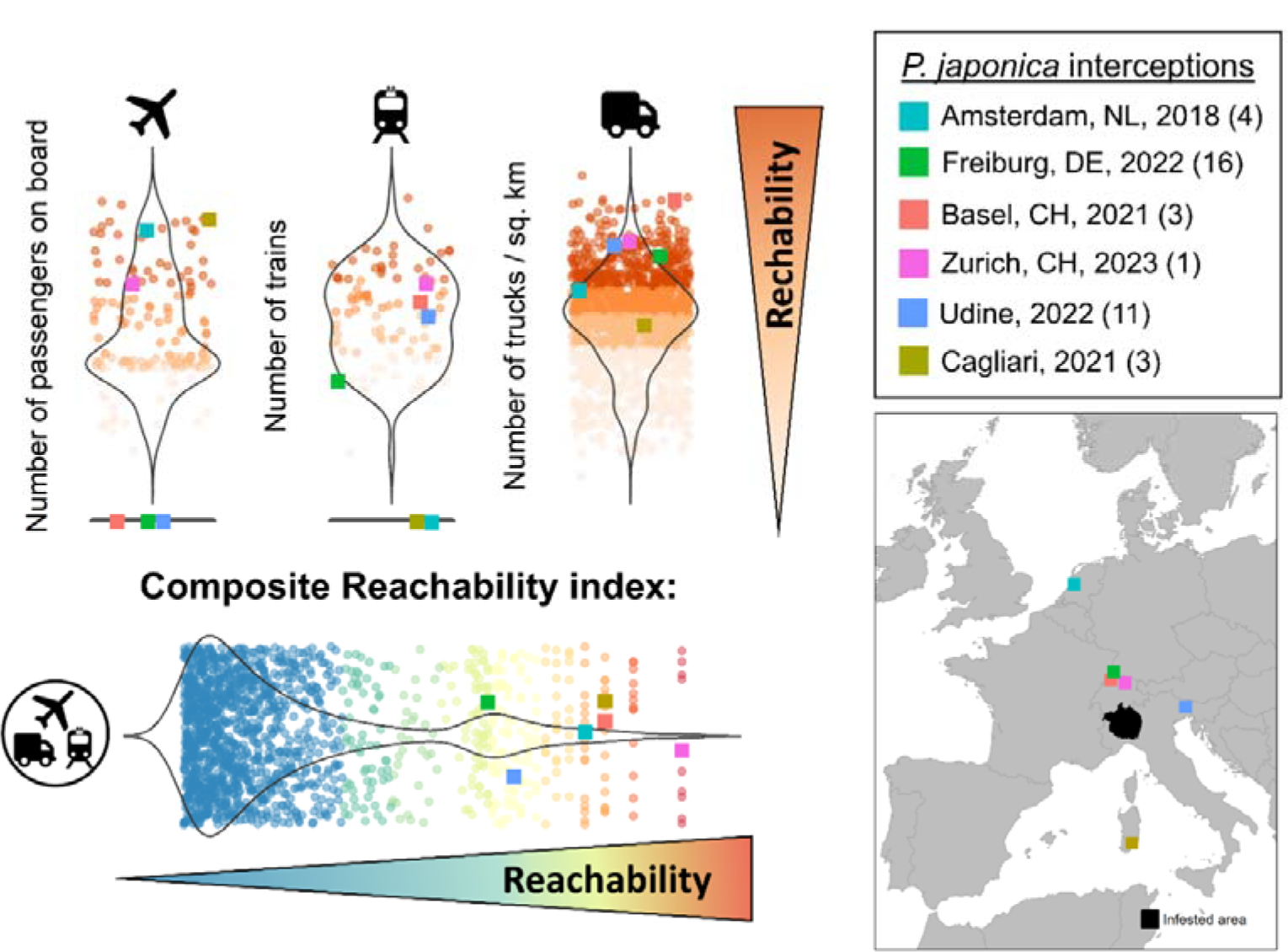
*P. japonica* interceptions made in Europe since 2018 and their position in relation to the distribution of reachability indices (air, rail, road, and composite). The number in brackets to the right of the interception site name indicates the group number assigned to the NUTS3 region by Pareto ranking, from most reachable (1) to least reachable (16).

The proposed framework provides a rapid response tool for decision-makers and phytosanitary services to anticipate the likelihood of hitchhiking pest entry on a continental scale. Informing risk-based surveillance strategies with likelihood of entry can drastically reduce surveillance efforts and promote early detection of invasive species (8). As data become available, further improvements may be achieved, for example by targeting commodity movements specifically identified as pest carriers (9), or by including other transport modes. Highly reachable regions could also be screened for the presence of sensitive host plants or favourable environmental conditions (10, 11). In conclusion, our framework highlights the need for local surveillance coupled with a transboundary strategy, involving official authorities and stakeholders, and adapted to the scale and means of spread of the pest under surveillance (12).

## Availability of data and materials

The datasets generated during the current study are available in the *French Research Government* repository, https://doi.org/10.57745/3WUVWJ.

## Funding information

This research was supported by the IPM-Popillia project, funded by the European Union Horizon 2020 research and innovation programme under grant agreement No 861852.

## Competing Interests

The authors have no relevant financial or non-financial interests to disclose.

## Ethical approval

This study did not require ethics approval.

## Notes

https://doi.org/10.57745/3WUVWJ

### Competing Interest Statement

The authors have declared no competing interest.

## References

1. P. E. Hulme, Trade, transport and trouble: managing invasive species pathways in an era of globalization. Journal of Applied Ecology 46, 10–18 (2009).

2. F. Essl, et al., Socioeconomic legacy yields an invasion debt. Proceedings of the National Academy of Sciences 108, 203–207 (2011).

3. P. Gotta, et al., Popillia japonica – Italian outbreak management. Frontiers in Insect Science 3 (2023).

4. Commission Delegated Regulation (EU) 2019/1702 of 1 August 2019, supplementing Regulation (EU) 2016/2031 of the European Parliament and of the Council by establishing the list of priority pests. [2019] L 260/8.

5. S. Poggi, L. Borner, J. Roche, C. Tayeh, D. Martinetti, “Biological invasion of the Japanese beetle in Continental Europe at a glance” (10.57745/R18NGL, Recherche Data Gouv, V3, 2022).

6. D. Speth, V. Sauter, P. Plötz, T. Signer, Synthetic European road freight transport flow data. Data in Brief 40, 107786 (2022).

7. R Core Team, R: A Language and Environment for Statistical Computing (R Foundation for Statistical Computing, 2021).

8. S. Parnell, T. R. Gottwald, T. Riley, F. van den Bosch, A generic risk-based surveying method for invading plant pathogens. Ecological Applications 24, 779–790 (2014).

9. G. Fenn-Moltu, et al., Alien insect dispersal mediated by the global movement of commodities. Ecological Applications 33, e2721 (2023).

10. A. J. Tatem, S. I. Hay, D. J. Rogers, Global traffic and disease vector dispersal. Proceedings of the National Academy of Sciences 103, 6242–6247 (2006).

11. L. Borner, D. Martinetti, S. Poggi, A new chapter of the Japanese beetle invasion saga: predicting suitability from long-invaded areas to inform surveillance strategies in Europe. Entomologia Generalis, in press (2023) 10.1127/entomologia/2023/2073.

12. A. Radici, D. Martinetti, D. Bevacqua, Global benefits and domestic costs of a cooperative surveillance strategy to control transboundary crop pathogens. PLANTS, PEOPLE, PLANET, 1–10 (2023).

