## Supplementary information - extended methods for "A hitchhiker’s guide to Europe: mapping human-mediated dispersal of the invasive Japanese beetle"

*Author names* : *Leyli Borner*[1], *Davide Martinetti*[2], *Sylvain Poggi*[1].

*Affiliations :*

*1. INRAE, Institut Agro, Univ Rennes, IGEPP, 35653, Le Rheu, France*
*2. INRAE, UR 546 BioSP, Avignon, France*

### Supplementary Information - extended methods

Data processing and analyses were performed using R version 4.2.1 (1).

For the airport analysis we used two databases: Eurostat - Detailed air passenger transport by reporting country and routes & The World Bank - Global airports. The Eurostat data were extracted for years 2010 to 2019 from May to August. The World Bank provides number of passengers on connecting flights between two airports for the year 2019. Some major airports were missing from the Eurostat database and were present in the World Bank data. We checked for correlation in the number of passengers for airports shared between the two databases ($R=0.95$, $p<0.001$, Pearson correlation). We fitted a GAM model with Poisson distribution on a training dataset of shared airports with World Bank number of passengers as explanatory variable and Eurostat number of passengers as the response variable (k=4, family=”poisson”, gam function of mgcv package 1.8-42 (2)). This model explained 84.6% of deviance found in data with a R-sq.(adj) of 0.891. We used this model to predict Eurostat number of passengers using World Bank number of passengers where data was missing from Eurostat.

For the train analysis, there were several steps including queries made to Deutsche Bahn Transport Rest API V5. For these queries, we used the httr2 package 0.2.2 (3).

1. We collected spatial coordinates of all train stations contained within the infested area. For this we used the EuroGlobalMap 2022 dataset (EGM 2022.2, © Institut national de l’information géographique et forestière (IGN-F), © EuroGeographics).
2. We used the “GET /stops/nearby” query to Deutsche Bahn Transport Rest API V5, using XY coordinates retrieved at the first step as arguments, retrieving the first/closest result within a 500 meters radius from given coordinate (‘&results=1&distance=500').
3. We retrieved trip ids for trains departing from the train stations within the infested zone, retrieved at step 2, from ('2022-05-01') to ('2022-08-31') (adults

emergence season). We used the “GET /stops/:id/departures” query, using IDs of train stations retrieved at step 2, and all departures at each date, for 1440 minutes (24h), retrieving 1000 results (departures were estimated to be less than 1000 per day) only retrieving results for trains. (duration=1440&when=', date, '&results=1000&remarks=false&bus=false&ferry=false&subway=false&tram =false&taxi=false')). We stored tripID and line names for each departures.

4. For each tripID retrieved at step 3, we retrieved all train stations were the trains stopped on this trip. We used the “GET /trips/:id” query, using the ID of each trip and the line name (stopovers=true&remarks=false&polyline=false&language=en')). As the final dataset we stored the id of trip, information on the station of departure, destination of the train, on the train line information. For each stop on the train paths we retrieved the station id, station name and coordinates, as well as the time of arrival and the time of departure.
5. Finally, we removed any duplicated data retrieved. We mapped all train stations and differed those that are within the infested area and those that are reached from the infested area. We removed the parts of trips that started outside of the infested area. For each train stations reached, we calculated the total number of trains reaching these stations by counting the number of unique trip ids at these stations.

For the truck analysis, we selected NUTS-3 from Speth et al. (4), which were either completely or partially covered by the infested area, as our ID_origin_region. As there was a perfect correlation between tons and trucks variables ($R=1$, $p<0.001$), and to be consistent with the train analysis, we decided to use the number of trucks variable (Traffic_flow_trucks_2019). For each destination region (ID_destination_region), we summed the number of trucks (Traffic_flow_trucks_2019). Finally, for each destination region, we weighted the total number of trucks arriving from the infested area, by the area of the destination region (in $km^2$).

The Pareto front analysis was carried out using rPref package 1.4.0 (5). We used the psel function with a complex preference function composing the preference for a Skyline query (also known as Pareto frontier) (p <- high(Traffic_flow_trucks_2019_p) * high(sumpassengers_EUROSTAT) * high (number_trains)); with a top value of number of 1449 that is the number of NUTS-3 reached sites (k-best tuples of the data set are returned).

NUTS-3 sites were chosen as the spatial resolution for the composite index of connectivity as it is the coarsest spatial resolution among transport modes. For trains and planes, the number of trains and number of passengers on board were summed among all train stations and airports contained within each NUTS-3 site.

### References


1. R Core Team, *R: A Language and Environment for Statistical Computing* (R Foundation for Statistical Computing, 2021).

2. S. N. Wood, Fast stable restricted maximum likelihood and marginal likelihood estimation of semiparametric generalized linear models. *Journal of the Royal Statistical Society: Series B (Statistical Methodology)* **73**, 3–36 (2011).

3. H. Wickham, *httr2: Perform HTTP Requests and Process the Responses* (2022).

4. D. Speth, V. Sauter, P. Plötz, T. Signer, Synthetic European road freight transport flow data. *Data in Brief* **40**, 107786 (2022).

5. P. Roocks, Computing Pareto Frontiers and Database Preferences with the rPref Package. *The R Journal* **8**, 393–404 (2016).
